# The genetic basis of resistance to pathogens in rainbow trout: a meta-QTL analysis

**DOI:** 10.64898/2026.02.13.705103

**Authors:** R. Rodríguez-Vázquez, A. M. Karami, D. Robledo, K. Buchmann

## Abstract

Rainbow trout is affected by a broad range of pathogens causing large economic losses and animal welfare concerns. Marker-assisted selection can significantly enhance resistance to pathogens in a few generations, and to this end many studies have focused on identifying quantitative trait loci (QTLs) for resistance traits. The integration of accumulated genetic resources provides an opportunity to uncover important genetic variation and candidate genes crucially involved in rainbow trout immunity. Here, we present a comprehensive meta-QTL (MQTL) analysis based on the integration of 145 QTLs related to pathogen resistance. These QTLs were refined into 26 MQTLs, of which 15 were validated by genome-wide association studies (GWAS). The average confidence interval (CI) of these MQTLs was reduced by 2.03-fold compared to the initial QTL, improving mapping precision. Integration of GWAS results revealed regions along the rainbow trout genome pivotal for pathogen resistance, and a major region in chromosome 3, which could be used in marker-assisted selection. Further, among the validated MQTLs we identified a subset of high-confidence MQTLs, based on those supported by at least three initial QTL from more than two independent studies, with a percentage of variance explained greater than 8% and a LOD score higher than three. Gene annotation identified 11 unique candidate genes within these high-confidence MQTLs involved in immune pathways, encoding proteins involved in the regulation of immune responses, signalling pathways, receptor activity, and direct immune effector production. The MQTLs and candidate genes identified are valuable resources for advancing molecular breeding and unravelling the genetic basis of pathogen resistance in rainbow trout.

## 1. INTRODUCTION

Rainbow trout (*Oncorhynchus mykiss*) is one of the most extensively farmed freshwater fish species, with a global production reaching 900 thousand tonnes per year (FAO, 2024). However, the intensification of aquaculture systems has facilitated the spread of pathogens, provoking severe challenges to fish health and welfare, environmental sustainability and economic viability (Houston et al., 2020; Shinn et al., 2015). Additionally, in the current context of climate change rising water temperatures further facilitate pathogen proliferation and spread (Fraslin et al., 2022). Rainbow trout is particularly vulnerable to disease outbreaks, such as Bacterial Cold Water Disease caused by *Flavobacterium psychrophilum*, furunculosis elicited by *Aeromonas salmonicida*, vibriosis due to *Vibrio anguillarum*, enteric redmouth disease caused by *Yersinia ruckeri* as well as viral haemorrhagic septicemia and white spot disease caused by *Ichthyophthirius multifiliis,* often resulting in substantial economic losses for producers (Campbell et al., 2014). Diseases in aquaculture are causing up to US$6 billion losses every year (Akazawa et al., 2010), and consequently, pathogen resistance has emerged as a critical trait to reduce the economic impact of disease outbreaks and support the sustainability of intensified production systems (Fraslin et al., 2018; Huang et al., 2024).

Resistance to infectious diseases in rainbow trout is known to be heritable and often associated with quantitative trait loci (QTLs) (Gleeson et al., 2000; Karami et al., 2020; Wiens et al., 2013). QTL mapping has facilitated the identification of genomic regions associated with resistance traits, accelerating marker-assisted selection (MAS) and the development of selective breeding programs (Buchmann, 2022). Over recent years, several QTLs conferring relative innate resistance towards pathogens such as *F. psychrophilum* (Mathiessen et al., 2023; Vallejo et al., 2014), *viral haemorrhagic septicemia virus* (Verrier et al., 2013), *V. anguillarum* (Karami et al., 2020) and *Ichthyophthirius multifiliis* (Jaafar et al., 2020) have been reported. However, the practical application of these QTLs in breeding programs has encountered challenges. A major issue is the limited overlap between QTL regions identified in independent studies, even when targeting resistance to the same pathogen. These discrepancies may be attributable to differences in experimental design, population genetic backgrounds, environmental conditions during infection or the statistical power and marker density of the mapping approach (Sharma et al., 2024; Yang et al., 2021). Moreover, early QTL studies with sparse marker coverage produced broad genomic intervals, complicating the identification of causal variants, especially in genomic regions with suppressed recombination or structural complexity (Calboli et al., 2022; Campbell et al., 2014; Villoutreix et al., 2021).

Meta-QTL (MQTL) analysis allows the integration of QTL data from multiple studies and the refinement of the position of loci associated with a given trait (Arcade et al., 2004; Goffinet and Gerber, 2000; Veyrieras et al., 2007). By aggregating and examining data from multiple studies, MQTL analysis enhances QTL localization through non-parametric models that are less susceptible to the biases and variability present in individual studies. This integration narrows the confidence intervals of QTLs derived from individual analyses, resulting in more stable, uniform and trustworthy consensus MQTL regions (Goffinet and Gerber, 2000; Khojasteh et al., 2024; Welcker et al., 2011). This refinement improves genetic resolution, facilitates the identification of robust markers for MAS and supports the discovery of candidate genes (Khahani et al., 2019; Tanin et al., 2022). While this approach has been widely and successfully applied in plant genetics (Bilgrami et al., 2023; Khahani et al., 2019; Khojasteh et al., 2024; Yang et al., 2021), its application in animal breeding remains limited, with only recent studies reported in pigs (Gonçalves da Silva et al., 2025) and buffalo (Sharma et al., 2021) for reproductive traits (Sharma et al., 2021). To date, no MQTL studies have been conducted in fish species for pathogen resistance. However, a recent preprint has reported meta-QTL analysis in rainbow trout for growth and fillet quality traits (Shakeri et al., 2025), highlighting the expanding application of these approaches. Thus, this study represents a novel application of MQTL analysis to pathogen resistance traits in aquaculture species, potentially providing valuable genetic insights.

In recent decades, advances in DNA sequencing technologies and high-throughput genotyping platforms have enabled genome-wide association studies (GWAS), which investigate the relationship between genomic variation at genome-scale and phenotypic traits. GWAS methodologies identify genetic markers significantly associated with target phenotypes, serving as potential QTLs and providing high-resolution mapping, as shown in *O. mykiss* for resistance to various pathogens (Calboli et al., 2022; Palti et al., 2024; Silva et al., 2019; Vallejo et al., 2019). Notably, several major QTLs identified through GWAS for resistance to *V. anguillarum, F. psychrophilum* and *I. multifilis* have been independently validated in subsequent studies (Buchmann et al. 2022; Karami et al. 2023; 2025; Mathiessen et al. 2023), reinforcing their biological relevance. These regions can be further explored to identify candidate genes involved in the genetic mechanisms underlying disease resistance. While GWAS offers high-resolution localization of trait-associated loci, traditional QTL mapping remains important for understanding the genetic architecture of complex traits. QTL mapping can detect genomic regions or alleles with moderate to large effects that may be omitted by GWAS due to limited statistical power in smaller cohorts or population-specific variation. The integration of QTLs identified through linkage mapping with those detected by GWAS provides complementary evidence, enhancing confidence in the robustness and biological relevance of the identified loci (Chen et al., 2018; Liu et al., 2025). MQTL-GWAS validated analysis is a powerful framework to synthesize data from both mapping approaches, refine the positions of consensus QTLs, and prioritize stable candidate regions and genes for functional studies and marker-assisted selection.

The objective of this study is to perform a comprehensive meta-QTL analysis of published QTLs related to pathogen resistance in rainbow trout and to integrate supporting evidence from recent GWAS studies. By integrating these complementary data sources, we aim to identify consensus genomic regions and key candidate genes underlying resistance traits in *O. mykiss*. The outcomes of this work will provide new insights into the genetic architecture of pathogen resistance in rainbow trout and establish a robust basis for the identification of important QTLs and candidate genes for use in breeding programs dedicated to disease-resilient aquaculture.

## 2. Material and methods

### 2.1 Collection of linkage mapping QTL studies for pathogen resistance

A comprehensive literature search was conducted for articles reporting QTLs associated with resistance to pathogens in *Oncorhynchus mykis*s. The following databases were used to collect the original research articles: Google Scholar (https://scholar.google.es/schhp?hl=es&as_sdt=2005&sciodt=0,5), Web of Science (https://www.webofscience.com/wos/woscc/basic-search), Scopus (https://www.scopus.com/search/form.uri?display=basic&zone=header&origin=AuthorProfile#basic) and NCBI (https://www.ncbi.nlm.nih.gov/). Search terms included combinations such as (“rainbow trout” OR “*Oncorhynchus mykiss*”) AND (“QTL” OR “quantitative trait loci”) AND (“resistance” OR “survival” OR “mortality”) AND (“pathogen” OR “disease” OR “infection” OR specific pathogens). Each QTL mapping study was meticulously examined from each independent study to extract the following data: 1) pathogen; 2) disease; 3) traits associated with pathogen/disease resistance; 4) mapping population (e.g. F2 generations, doubled haploids (DH), backcrosses (BC)) and their parental lines; 5) population size; 6) chromosome number; 7) the position of the QTLs and flanking markers; 8) logarithm of odds (LOD) values; 9) variance explained by the individual QTLs (PVE or r^2^). Each QTL was considered as an independent entity, even if they were identified in multiple environments or genetic backgrounds. When QTL studies reported likelihood ratio statistics (LRS) scores, the values were converted to LOD scores using the following formula (Liu, 1998):

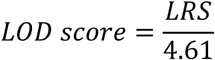

For the very few QTLs that did not have LOD/LRS and R^2^/PVE values, these were assumed as 3 and 10%, respectively, following common practice (Yang et al., 2021). The basic information of these QTLs is summarized in Table S1 and S5.

### 2.2. Development of a comprehensive consensus map

Six genetic maps, each containing multiple markers and commonly utilized in various QTL mapping studies, were employed to construct a reference genetic map “Rainbow trout Consensus map_2025” (Table S2). The maps incorporated several common markers situated at various genetic positions, which were integrated during the development of the consensus maps by the R package “LPmerge” (Endelman and Plomion, 2014). The fundamental principle of this methodology is to categorize co-segregating markers into “bins”, thereby ensuring that the consensus map retains marker order consistency with the individual linkage maps to the greatest extent possible. LPmerge achieved this by removing the smallest “bins” from each map. LPmerge effectively resolves marker order discrepancy across different maps, as described by Khojasteh et al., 2024 (Table S3).

### 2.3 Projection of QTLs

The novel consensus genetic map of rainbow trout was employed to project QTL positions by using the homothetic method described by Chardon et al., 2004 and implemented in the software Biomercator V4.2.3, with the QTLProj command. This approach is based on the principle of a scaling rule that is derived from the relative positions of the flanking markers in the original maps and their corresponding locations in the consensus map (Arcade et al., 2004; Sosnowski et al., 2012). To carry out a QTL projection, the software requires a set of distinct parameters, including chromosome name, LOD score, PVE value, genetic peak position and CIs of the QTL, the trait linked with the QTL, and the size of the mapping population that was used to identify the QTL.

When original publications did not report the 95% CIs for a specific QTL we adapted population-specific formulas (Darvasi & Seller, 1997; Guo et al., 2006; Venske et al., 2019), which were used to calculate the 95% CIs as follows:

- For F_2_ and Backcross populations: 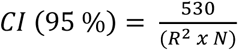
- For Double Haploid (DH) population: 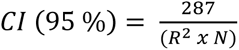

where R^2^ is defined as the phenotypic variance explained by the individual QTL and *N* is population size. The QTLs which could not be mapped onto the consensus map were discarded from the meta-analysis. Nevertheless, those QTLs were still collected and characterized in a comprehensive catalogue, and the details of these initial QTLs are provided in Table S5.

## 2.4 QTL meta-analysis approach

QTL meta-analysis was performed separately for each rainbow trout chromosome employing algorithms available in BioMercator software v4.2.3. Two distinct methods were applied depending on the number of projected QTLs on each chromosome. For chromosomes with 10 or fewer projected QTLs, the method described by Goffinet & Gerber, 2000 was applied. In this method, Biomercator evaluates five alternative MQTL models, each associated with a distinct Akaike Information Criterion (AIC) value, to estimate the most probable MQTL configuration for that chromosome. The model with the lowest AIC was selected to define the MQTL position on the consensus map and any MQTL region represented by only a single QTL was discarded. For chromosomes displaying more than 10 projected QTLs, the method described by Veyrieras et al. (2007) was used. This procedure was comprised of two main steps: 1) clustering the projected QTLs on each chromosome using default parameters and 2) determining the optimal number of MQTLs based on five selection criteria: AIC, AICc, AIC3 (modified AICs), BIC (Bayesian Information Criterion), and AWE (Approximate Weight of Evidence). The best MQTL model was determined to be the one with the lowest values in at least three out of the five specific criteria. The selected optimal model was then used to determine the number of MQTLs per chromosome, their peak positions, and the 95% CIs for each MQTL. QTLs were clustered such that the peak of the original QTLs fell within the CI of the corresponding MQTL. MQTLs were designated based on chromosome number and their relative position (e.g. “MQTL1.1”, “MQTL1.2”). Visualization of QTL and MQTL were carried out in the *R* statistical environment using custom scripts (v4.4.2; R Core Team, 2024). Graphical representations, including circus plots illustrating chromosomal distribution and patterns of QTLs and MQTLs were generated with the R package circlize (V. 0.4.16) (Gu et al., 2014).

### 2.5 Identifying significant genomic regions based on GWAS studies

Genome-wide association study (GWAS) data reporting SNP markers (p < 0.05) and genomic locations associated with relevant traits were collected from studies published. Detailed information for these GWAS studies is provided in Table S6. Reported SNP peak signals from GWAS studies, originally mapped to different *O. mykiss* genome assemblies, were first standardized to the OmykA_1.1 reference genome (GCA_013265735.3) using local BLASTN program. Genomic regions containing SNPs from three or more signals and from at least two independent GWAS studies were identified and prioritized for integration with meta-QTL regions from linkage mapping studies (Section 2.6). This approach provided an aggregated and quantitative assessment of recurrent association genomic regions, reinforcing candidate locus characterization through the convergence of multiple independent evidence. The processing and analysis of the GWAS data, as well as the identification of genomic regions, were conducted using the R statistical environment. Data manipulation, grouping and summarization were performed with functions from the dplyr package (V. 1.1.4), while graphical visualization was carried out using ggplot2 (V. 3.5.1.).

### 2.6 Validation of predicted MQTL by GWAS studies

The MQTLs were mapped to the rainbow trout genome to obtain their physical position. The markers located at the flanking regions on both sides of each MQTL confidence interval were manually identified, and their sequences retrieved. These flanking sequences were then aligned to the rainbow trout reference genome (OmykA_1.1; GCA_013265735.3) using the local BLASTN program (minimum sequence identity of 95%) to determine the physical location of these markers. In cases where the genomic positions of the flanking markers could not be identified, the closest markers were used to assign MQTL genomic coordinates (Li et al., 2025).

To validate the predicted MQTLs the significant SNPs identified in GWAS studies were compared with the MQTL regions to assess collinearity and commonality between detected MQTL intervals and GWAS signals. This cross-validation allowed us to verify the accuracy and robustness of MQTLs by confirming their overlap with independently detected association signals.

### 2.7 Functional analysis and identification of candidate genes for disease resistance

The MQTLs validated with the reported genome-wide association studies were further analysed to obtain predicted genes between the flanking markers. For the functional analysis, only high-confidence MQTLs were selected according to the quality criteria adapted from Sharma et al., 2024 and Acuña-Galindo et al., 2015: i) contain at least three original QTLs; ii) have a LOD score ≥ 3; iii) PVE > 8%, iv) include QTLs reported in at least two independent studies and v) being GWAS validated.

To identify candidate genes in these high-confidence QTL regions, two complementary strategies were applied. The first strategy, adapted from Pal et al. (2021), was based on the physical interval defined by the flanking markers. MQTL having a flanking marker interval < 2 Mb were directly used for the exploration of genes, while for those MQTL having marker interval > 2 Mb, the search was restricted to a 2 Mb window centred on the calculated peak position, corresponding to ± 1Mb upstream and downstream. The peak position (bp) was determined by the formula:

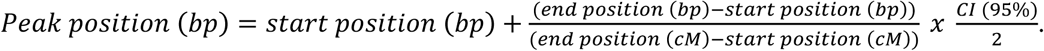

The second strategy focused on SNPs located within the validated-MQTL regions, a strategy adapted from Shakeri et al. (2025). All SNPs reported in the GWAS studies (Table S6) whose genomic positions fell within MQTL CI were identified. For each MQTL, these SNPs were extracted with their annotated genes from the rainbow trout reference genome, providing candidate genes with direct GWAS association evidence.

For both strategies, candidate genes were mined from the *Oncorhynchus mykiss* genome using the Ensembl BioMart tool and the rainbow trout genome annotation (v 2.62.1) (Durinck et al., 2005). Gene Ontology (GO) terms associated with these genes were also obtained and subsequently were filtered using AmiGO 2 (Carbon et al., 2009) and QuickGO (Binns et al., 2009) to retain only genes associated with immune or defence responses; genes associated with GO terms contained the keywords “immune system process” were considered candidate genes.

## 3. RESULTS

### 3.1 Distribution of the QTLs associated with pathogen resistance along the rainbow trout genome

A comprehensive review of relevant literature was conducted employing major databases. A total of 145 QTLs associated with pathogen resistance traits were compiled for subsequent meta-QTL analysis. The distribution of QTLs per chromosome exhibited variation, as is illustrated in Fig. 1A, with a minimum of one QTL observed on chromosome 1 and a maximum of 22 QTLs on chromosome 8 (Table S1). Moreover, as it is shown in Fig. 1B, the majority of the initial QTLs were derived from studies focusing on resistance to *F. psychrophilum*. The LOD values of initial QTLs ranged from 1.05 to 8.65, although the majority of QTLs (88.96% of the total) exhibited LOD scores between 2.0 to 3.0, and the average was 3.06 (Fig. 1C; Table S5). The percentage of PVE by individual QTL varied widely, from 1.72 % to 72%, with an overall mean of 15.38%. Approximately, 13.79% of QTLs had a PVE value less than 5%, and only a small fraction (7.58%) exhibited PVE values above 30%, which suggests the involvement of both major- and minor-QTLs in governing resistance traits (Fig. 1D; Table S5).

**Fig. 1.**
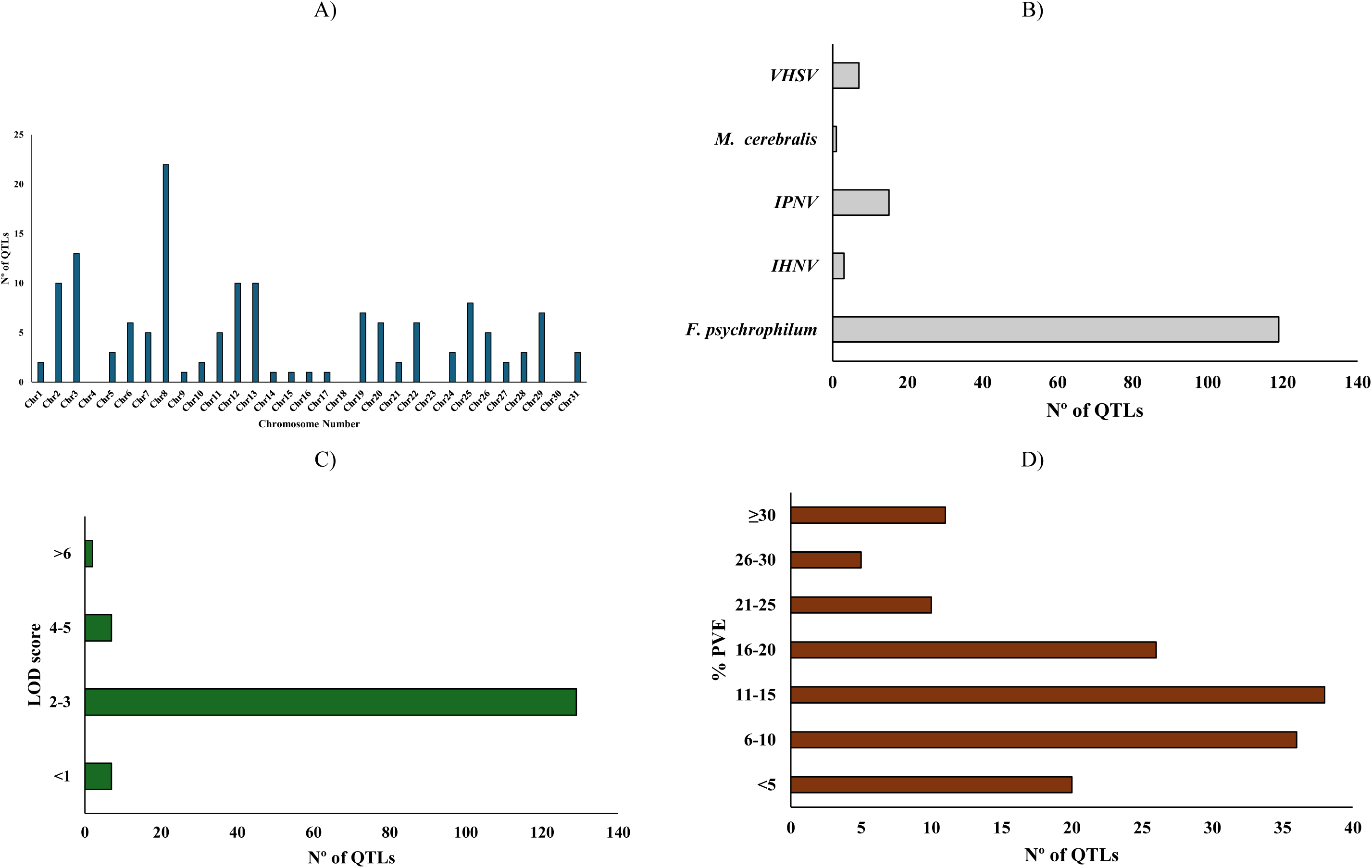
Summary of QTLs associates with pathogen resistance. A) Chromosome-wise distribution of initial QTLs; B) Distribution of initial QTL associated with pathogen resistance traits; C) LOD scores of initial QTLs. D) % PVE of the initial QTLs

### 3.2 Rainbow trout consensus map and QTL projection

The high-density “Rainbow trout Consensus map_2025” constructed in this study by integrating six genetic maps revealed substantial variation in marker distribution across chromosomes (Table S2). Marker density on individual chromosomes varied from 1.72 markers/cM on chromosome 24 to 4.27 markers/cM on chromosome 4, with a mean of 2.93 markers/cM across the genome (Fig. 2; Table S3 and S4). Chromosome lengths also varied notably, spanning from 99.0 cM for chromosome 28 to 265.6 cM for chromosome 3, with an average of 182.23 cM. The cumulative genetic map length of all chromosomes in the map was 5284.6 cM, spanned by 15,381 markers. On average, each chromosome contained approximately 530.4 mapped markers, with numbers ranging from a few hundred (237 on chromosome 26) to several hundred (763 on chromosome 12).

**Fig. 2.**
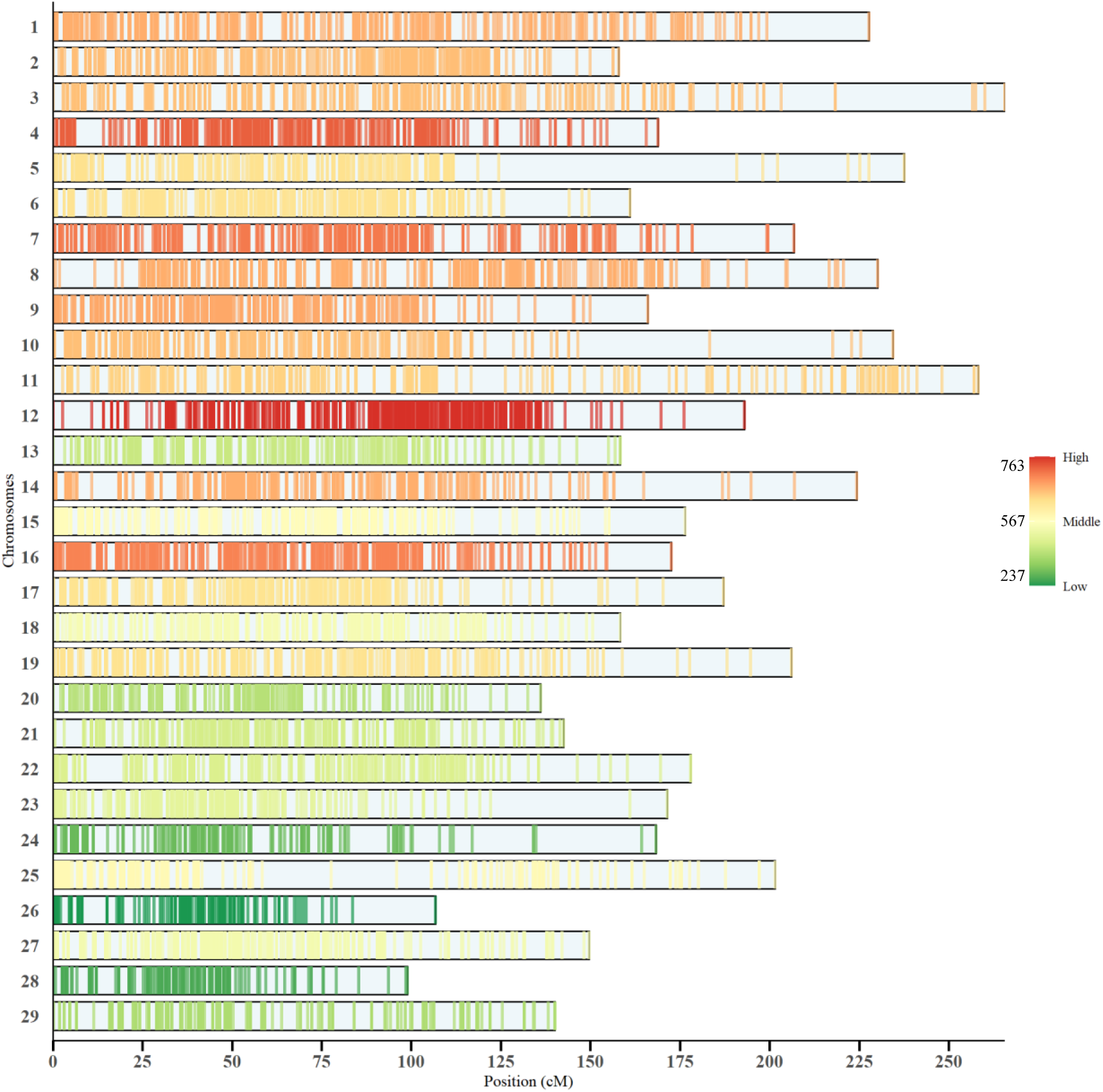
Marker distribution on the consensus genetic map used for meta-QTL analysis. Vertical lines represent individual markers. Marker lines coloured on a continuous scale by total marker density per chromosome: light green (low density, 237 markers), yellow-orange (middle density, 567 markers), dark-red (high density, 763 markers). Light blue backgrounds represent chromosomes without markers.

### 3.3 Meta-analysis of QTLs associated with pathogen resistance

In total, 145 initial QTLs were mapped to the consensus map, of which 100 QTLs (72.46% of the total) could be projected (Table S5). The remaining QTLs could not be projected due to either an insufficient number of shared markers between the consensus and initial genetics maps or large CIs associated with the initial QTLs (Sharma et al., 2024).

A total of 26 MQTLs were predicted for pathogen-resistance-related traits, of which 22 were associated exclusively with *F. psychrophilum*, while four MQTLs (MQTL 3.1, MQTL 12.1, MQTL 22 and MQTL-26) were associated with both *F. psychrophilum* and IPNV (Table 1). The 26 MQTL were distributed across different chromosomes (Fig. 3A) and involved different numbers of QTLs from independent studies in each MQTLs (Fig. 3B). Their characteristics and distributions of initial and projected QTLs and MQTLs illustrated in Fig. 3C, Fig. 4, Fig. S1 and Table S5. No MQTLs were predicted on chromosomes 4, 9, 10, 14, 15, 16, 17, 18, 19, 23, 27 and 28. The highest number of MQTL were located on chromosome 8 and 12, containing 4 and 3 MQTL, respectively (Fig. 3A, Fig. 4 and Table 1). Overall, 73.08% MQTLs were composed of three or more QTLs (Fig. 3B, Table 1). The PVE by individual MQTL varied from a minimum of 4.86% to a maximum of 25.5%, with a mean of 12.02%, while the LOD score varied from 1.68 to 4.58 with an average of 3.03, finally the 95% of confidence interval of predicted MQTLs ranged from 5.81 to 33 cM (Fig 4, Table 1). On average, the CI of MQTLs were significantly reduced by a factor of 2.03 compared to the initial QTLs. Significant differences in CI reduction were observed across rainbow trout chromosomes; the average CI was reduced by 3.04 and 3.01-fold less on chromosomes 6 and 8 respectively, followed by reductions of 2.80, 2.73 and 2.76 on chromosomes 5, 12 and 13 (Fig. 3C)

**Fig. 3.**
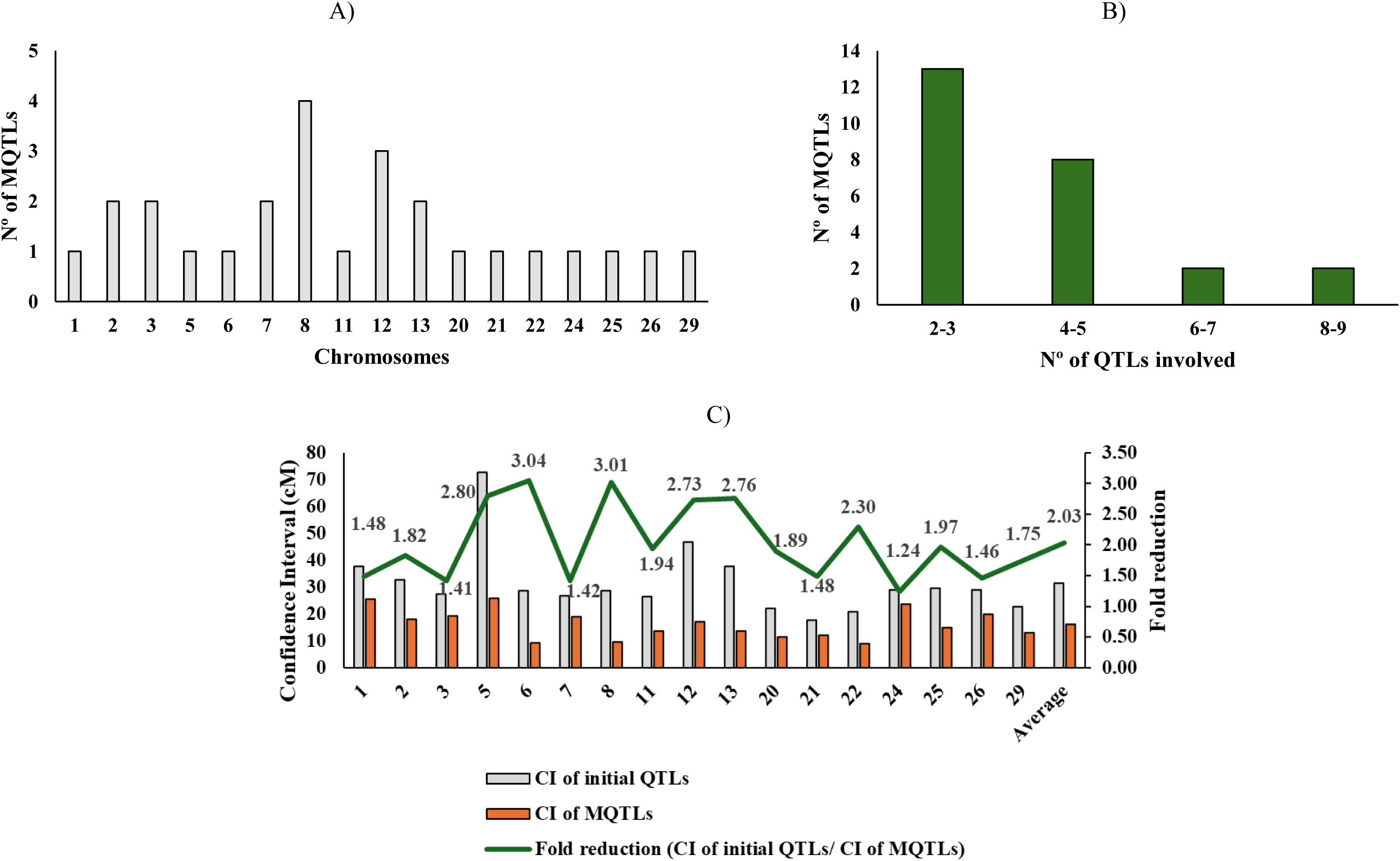
Basic information of MQTLs associated with pathogen resistance. A) Chromosome wide distribution of MQTLs. B) Number of QTLs involved in different MQTLs. C) Reduction of QTL confidence interval (CI, 95%) after meta-QTL analysis. The grey and orange bars represent the average CI length (cM) of original QTL used for projection and of the MQTL on chromosome, respectively, and the green line represents the fold change of the QTL CI length.

**Fig 4.**
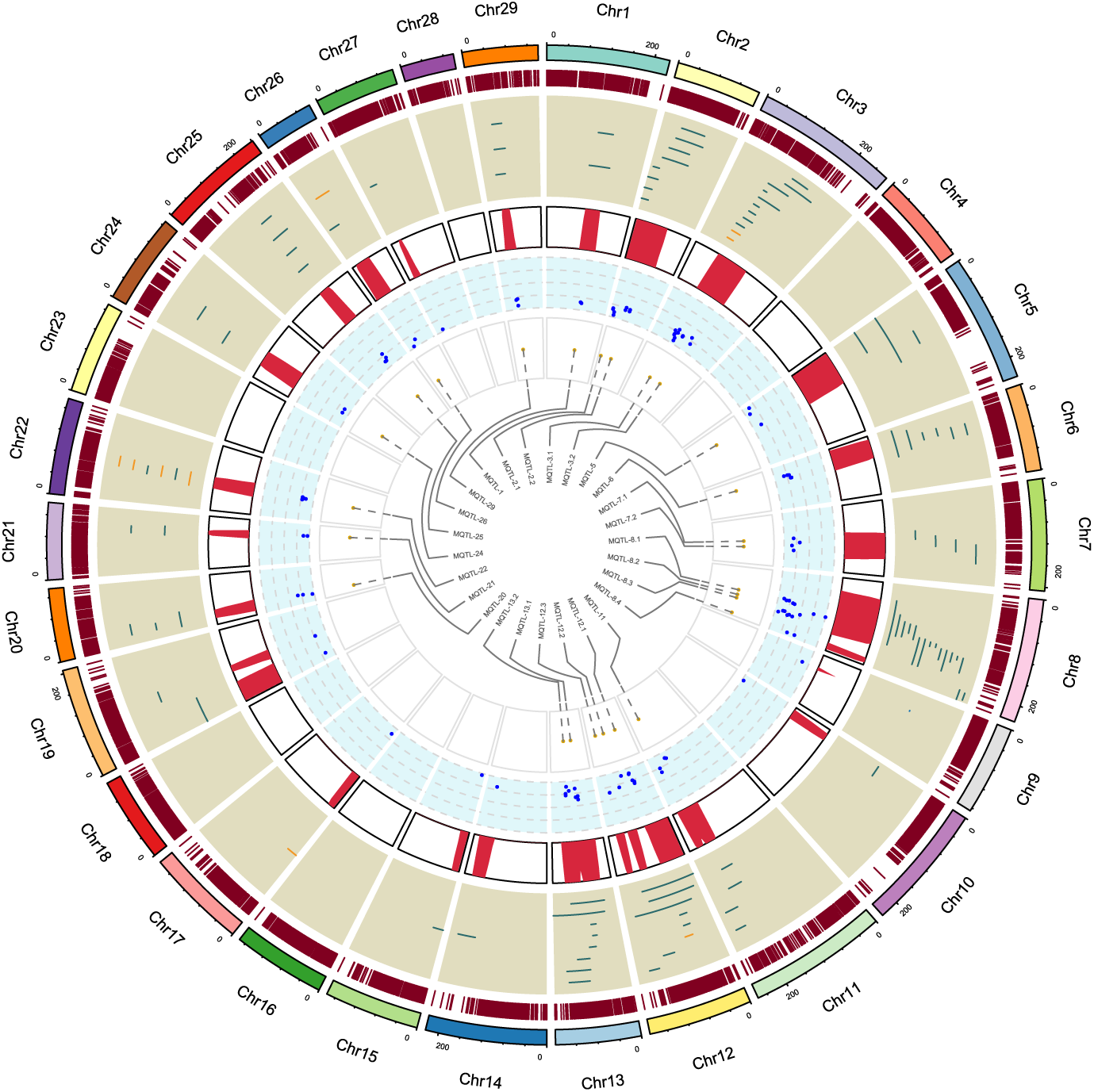
Circos diagram showing the genome-wide distribution of MQTLs. From the outermost to the innermost circle: Track 1 represents the chromosome positions (cM). Track 2 outlines the consensus map density, with red bands indicating marker density. Track 3 displays the distribution of projected QTLs used in this MQTL analysis (green for *F. psychrophilum* resistance, orange for IPNV resistance). The fourth and five tracks show density and PVE value of projected QTL. The innermost circle displays position of MQTLs on the consensus genetic map.

**Table 1.**
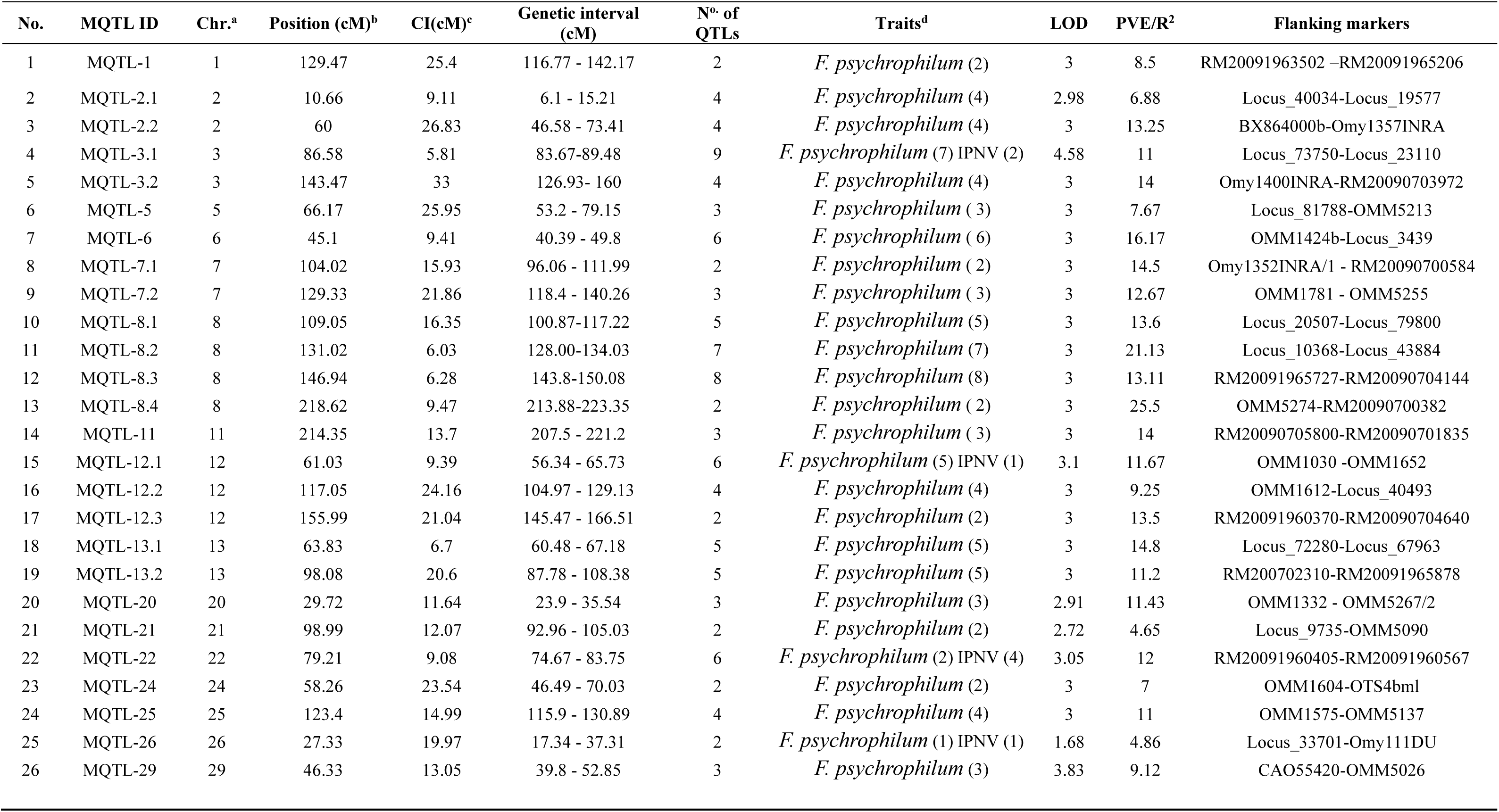
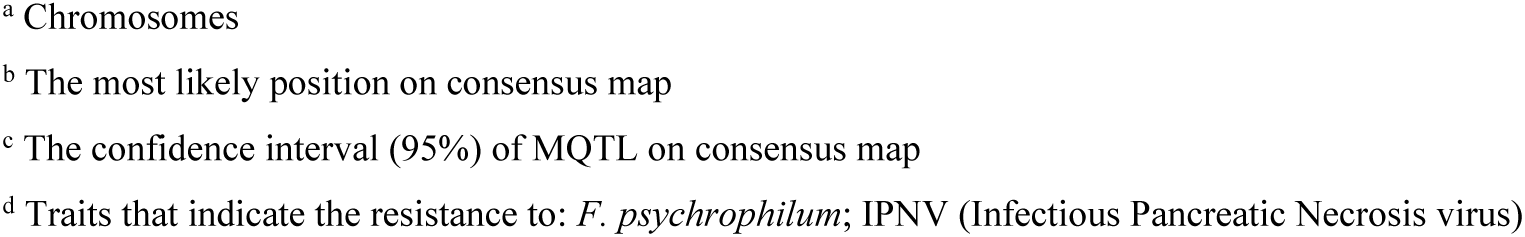
MQTLs associated with pathogen resistance in the present study.

| No. | MQTL ID | Chr.<sup>a</sup> | Position (cM)<sup>b</sup> | CI(cM)<sup>c</sup> | Genetic interval (cM) | N<sup>o</sup>. of QTLs | Traits<sup>d</sup> | LOD | PVE/R<sup>2</sup> | Flanking markers |
| --- | --- | --- | --- | --- | --- | --- | --- | --- | --- | --- |
| 1 | MQTL-1 | 1 | 129.47 | 25.4 | 116.77 - 142.17 | 2 | *F. psychrophilum* (2) | 3 | 8.5 | RM20091963502 –RM20091965206 |
| 2 | MQTL-2.1 | 2 | 10.66 | 9.11 | 6.1 - 15.21 | 4 | *F. psychrophilum* (4) | 2.98 | 6.88 | Locus_40034-Locus_19577 |
| 3 | MQTL-2.2 | 2 | 60 | 26.83 | 46.58 - 73.41 | 4 | *F. psychrophilum* (4) | 3 | 13.25 | BX864000b-Omy1357INRA |
| 4 | MQTL-3.1 | 3 | 86.58 | 5.81 | 83.67-89.48 | 9 | *F. psychrophilum* (7) IPNV (2) | 4.58 | 11 | Locus_73750-Locus_23110 |
| 5 | MQTL-3.2 | 3 | 143.47 | 33 | 126.93- 160 | 4 | *F. psychrophilum* (4) | 3 | 14 | Omy1400INRA-RM20090703972 |
| 6 | MQTL-5 | 5 | 66.17 | 25.95 | 53.2 - 79.15 | 3 | *F. psychrophilum* (3) | 3 | 7.67 | Locus_81788-OMM5213 |
| 7 | MQTL-6 | 6 | 45.1 | 9.41 | 40.39 - 49.8 | 6 | *F. psychrophilum* (6) | 3 | 16.17 | OMM1424b-Locus_3439 |
| 8 | MQTL-7.1 | 7 | 104.02 | 15.93 | 96.06 - 111.99 | 2 | *F. psychrophilum* (2) | 3 | 14.5 | Omy1352INRA/1 - RM20090700584 |
| 9 | MQTL-7.2 | 7 | 129.33 | 21.86 | 118.4 - 140.26 | 3 | *F. psychrophilum* (3) | 3 | 12.67 | OMM1781 - OMM5255 |
| 10 | MQTL-8.1 | 8 | 109.05 | 16.35 | 100.87-117.22 | 5 | *F. psychrophilum* (5) | 3 | 13.6 | Locus_20507-Locus_79800 |
| 11 | MQTL-8.2 | 8 | 131.02 | 6.03 | 128.00-134.03 | 7 | *F. psychrophilum* (7) | 3 | 21.13 | Locus_10368-Locus_43884 |
| 12 | MQTL-8.3 | 8 | 146.94 | 6.28 | 143.8-150.08 | 8 | *F. psychrophilum* (8) | 3 | 13.11 | RM20091965727-RM20090704144 |
| 13 | MQTL-8.4 | 8 | 218.62 | 9.47 | 213.88-223.35 | 2 | *F. psychrophilum* (2) | 3 | 25.5 | OMM5274-RM20090700382 |
| 14 | MQTL-11 | 11 | 214.35 | 13.7 | 207.5 - 221.2 | 3 | *F. psychrophilum* (3) | 3 | 14 | RM20090705800-RM20090701835 |
| 15 | MQTL-12.1 | 12 | 61.03 | 9.39 | 56.34 - 65.73 | 6 | *F. psychrophilum* (5) IPNV (1) | 3.1 | 11.67 | OMM1030 -OMM1652 |
| 16 | MQTL-12.2 | 12 | 117.05 | 24.16 | 104.97 - 129.13 | 4 | *F. psychrophilum* (4) | 3 | 9.25 | OMM1612-Locus_40493 |
| 17 | MQTL-12.3 | 12 | 155.99 | 21.04 | 145.47 - 166.51 | 2 | *F. psychrophilum* (2) | 3 | 13.5 | RM20091960370-RM20090704640 |
| 18 | MQTL-13.1 | 13 | 63.83 | 6.7 | 60.48 - 67.18 | 5 | *F. psychrophilum* (5) | 3 | 14.8 | Locus_72280-Locus_67963 |
| 19 | MQTL-13.2 | 13 | 98.08 | 20.6 | 87.78 - 108.38 | 5 | *F. psychrophilum* (5) | 3 | 11.2 | RM200702310-RM20091965878 |
| 20 | MQTL-20 | 20 | 29.72 | 11.64 | 23.9 - 35.54 | 3 | *F. psychrophilum* (3) | 2.91 | 11.43 | OMM1332 - OMM5267/2 |
| 21 | MQTL-21 | 21 | 98.99 | 12.07 | 92.96 - 105.03 | 2 | *F. psychrophilum* (2) | 2.72 | 4.65 | Locus_9735-OMM5090 |
| 22 | MQTL-22 | 22 | 79.21 | 9.08 | 74.67 - 83.75 | 6 | *F. psychrophilum* (2) IPNV (4) | 3.05 | 12 | RM20091960405-RM20091960567 |
| 23 | MQTL-24 | 24 | 58.26 | 23.54 | 46.49 - 70.03 | 2 | *F. psychrophilum* (2) | 3 | 7 | OMM1604-OTS4bml |
| 24 | MQTL-25 | 25 | 123.4 | 14.99 | 115.9 - 130.89 | 4 | *F. psychrophilum* (4) | 3 | 11 | OMM1575-OMM5137 |
| 25 | MQTL-26 | 26 | 27.33 | 19.97 | 17.34 - 37.31 | 2 | *F. psychrophilum* (1) IPNV (1) | 1.68 | 4.86 | Locus_33701-Omy111DU |
| 26 | MQTL-29 | 29 | 46.33 | 13.05 | 39.8 - 52.85 | 3 | *F. psychrophilum* (3) | 3.83 | 9.12 | CAO55420-OMM5026 |

- <sup>a</sup> Chromosomes - <sup>b</sup> The most likely position on consensus map - <sup>c</sup> The confidence interval (95%) of MQTL on consensus map - <sup>d</sup> Traits that indicate the resistance to: *F. psychrophilum*; IPNV (Infectious Pancreatic Necrosis virus)

### 3.4 Validating the MQTLs by GWAS studies

#### 3.4.1. Identifying significant genomic regions based on GWAS studies

The GWAS datasets encompassed all chromosomes, ranging from one significant SNP on Chr18 to multiple SNPs on Chr5, 16 and 17 (Table S6). Genomic regions containing ≥ 3 significant SNPs and supported by more than two independent studies were prioritised (Fig. 5, Table S7). These were identified on Chr1, Chr3, Chr6, Chr11, Chr 17, Chr21, Chr31. These regions represent candidate loci where multiple significant SNPs co-localized, highlighting potential functional genomic regions influencing diseases resistance (Fig. 5, Table S7). Noteworthy, validated regions at Chr17 (75–76 Mb) to 3 pathogens; Chr1 (12-13 Mb), Chr3 (51-52 Mb and 60-61 Mb), Chr11 (42-43 Mb) and Chr21 (22–23 Mb and 48–49 Mb) were associated with 2 pathogens respectively, while the remaining Chr6 (25-26 Mb) and Chr31 (20-21 Mb and 25–26 Mb) regions were pathogen specific (Fig. 5, Table S7).

**Fig 5.**
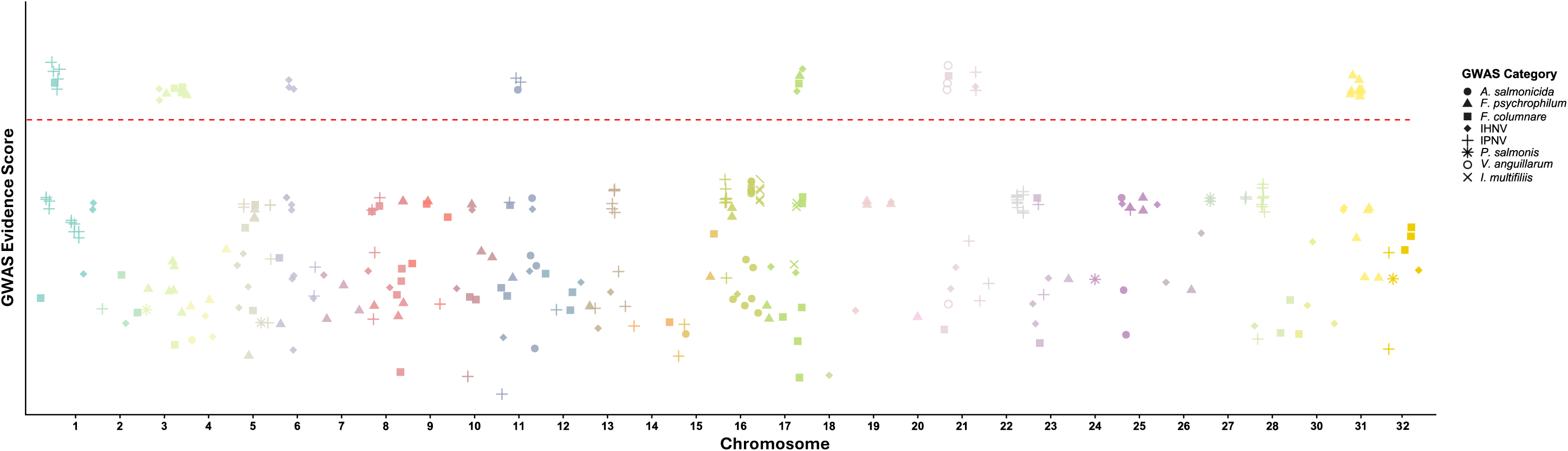
Manhattan plot representation of GWAS data across the rainbow trout genome. Each point corresponds to a SNP reported in Table S6. Shapes correspond to pathogen categories and colours denote chromosomes. The dashed horizontal line indicates significant genomic regions, defined as 1-Mb clusters containing more than three significant SNPs and supported by at least 2 independent studies (Table S7)

#### 3.4.2 Verification of MQTL with GWAS

Following the identification of significant resistance-associated regions, we further evaluated the reliability of MQTL analysis by overlaying them with the GWAS results (Table S6). Using the flanking markers, the physical positions of 23 MQTL were mapped to the rainbow trout genome (Fig. 6). However, the exact physical positions of three MQTLs (MQTL 8.4, MQTL 12.3 and MQTL 29) could not be ascertained due to unavailability of nucleotide sequences of the markers flanking these MQTL (Table S8). The physical intervals of these 23 MQTLs varied from 0.74 Mb to 38.9 Mb (Fig. 6, Table S8).

**Fig. 6.**
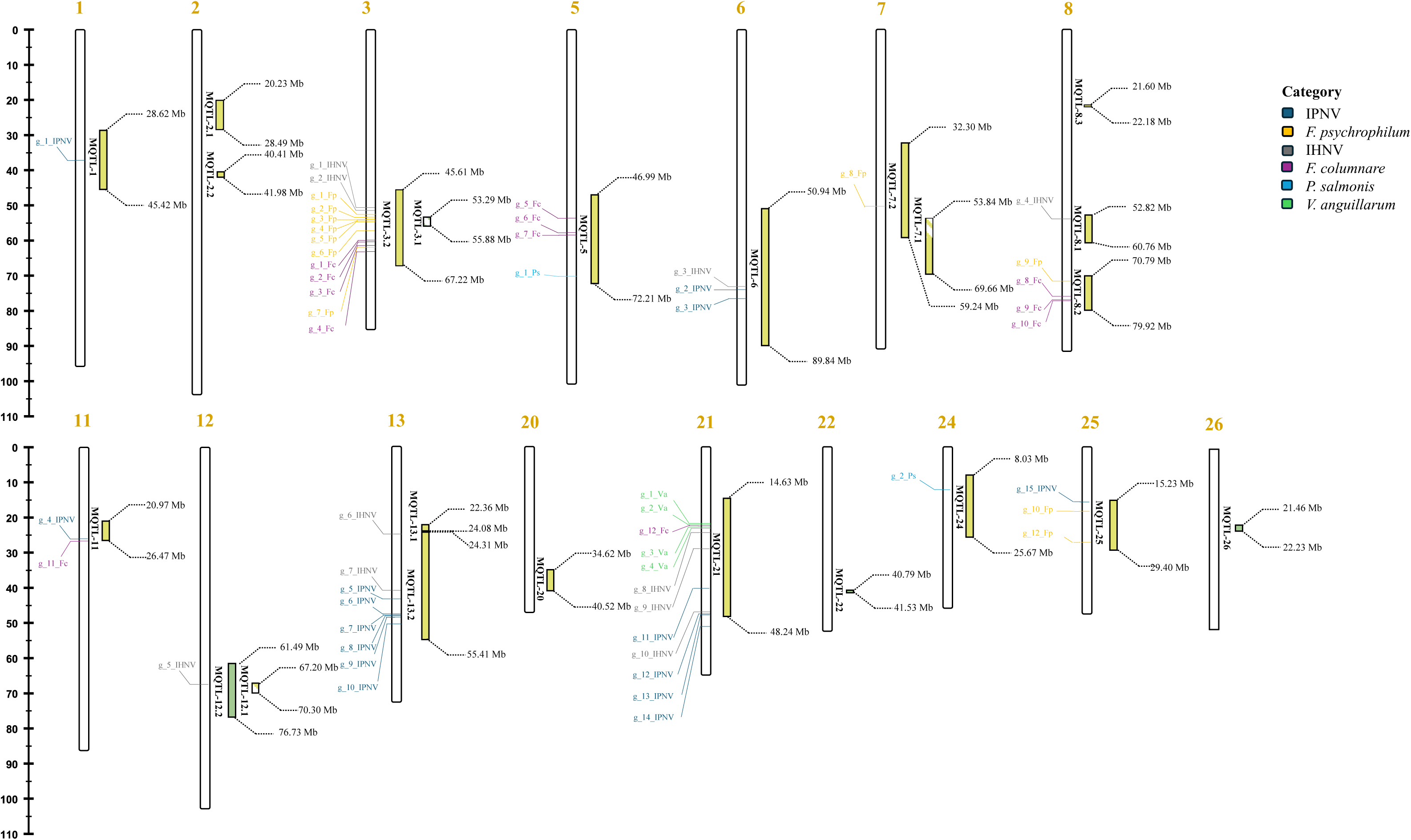
Distribution of MQTLs (Table 1) and GWAS loci (Table S6) on chromosomes associated with pathogen resistance. The chromosome map shows the GWAS loci on the left of the chromosome and MQTLs on the right, identified in this study and their co-location in a physical map. Light green indicates MQTLs formed by only one type of pathogen resistance QTL, while dark green indicates MQTLs formed by two different types of pathogen resistance QTLs. The striped areas indicate the overlap of two MQTLs. The axis on the left represents the physical distance (Mb). Detailed information is available in Table S8.

The physical coordinates of these MQTL identified in our study were compared to data reported in 16 previous GWAS that comprised a total of 297 SNPs for pathogen resistance (Table S6). Among these 23 MQTLs identified in physical map, a total of 15 MQTLs were found to overlap with GWAS studies (Table S8). Several MQTLs were found to be co-localized with significant SNPs from multiple GWAS studies, in fact 60 % of these MQTLs were co-localized with regions identified in two or more different GWAS. In addition, MQTL 3.2 and 21 co-localized with at least six independently reported GWAS (Table S8).

### 3.5 High-confidence MQTL and candidate gene mining

To enhance the reliability of the validated MQTLs we further refined them, leading to the identification of regions referred to as high-confidence MQTLs (hcMQTLs). In general, each hcMQTL cluster included a minimum of three initial QTLs, from more than two independent studies, with a PVE value greater than 8% and a LOD score greater than three. These regions were distributed across 7 chromosomes, obtaining the following 8 hcMQTLs (Table S8). To explore the potential functions of hcMQTL regions and prioritize genes associated with resistance, two complementary approaches were applied. The first focused on genes located in flanking markers or ±1 Mb around peak positions, and the second genes overlapping significant SNPs in these regions. This dual approach allowed us to identify a broad catalogue of genes with possible involvement in immune processes (Table 2, S9, S10). The interval-based method identified 201 genes (Table S9), and the SNP-based method identified 264 genes (Table S10).

**Table 2.**
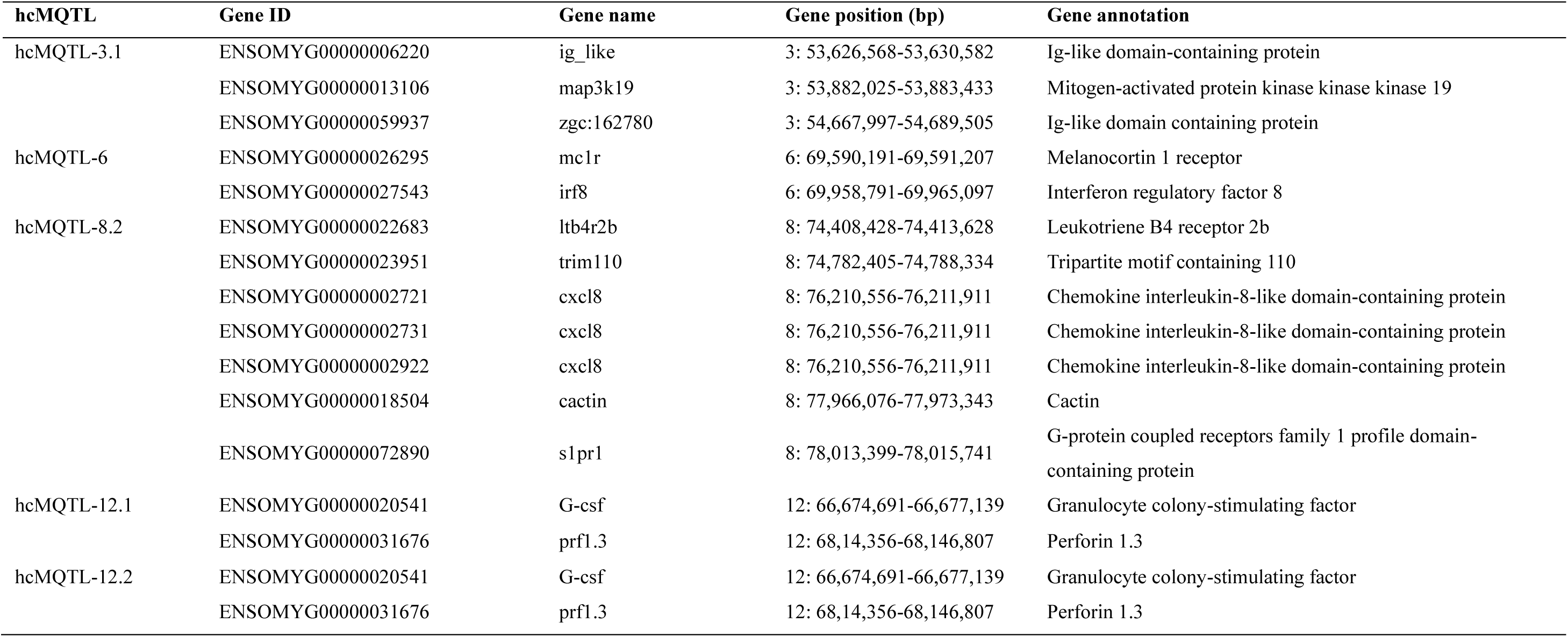
Candidate genes associated with immune system highlighted in hcMQTLs.

Filtering for immune-related GO terms revealed relevant candidates in five hcMQTLs. HcMQTL-3.1 included two Ig-like domain-containing protein and *map3k19*; hcMQTL-6.1 (Chr6: 69.5–69.9 Mb) contained *irf8* and *mc1r*; hcMQTL-8.2 (Chr8: 74.0–78.0 Mb) harboured several paralogs of *cxcl8, ltb4r2b*, *trim110*, *cactin*, and *s1pr1* and hcMQTL-12.1/12.2 (Chr12: 67.7–70.1 Mb) showed *prf1.3* and *g-csf* (Table 2).

## 4. DISCUSSION

Diseases caused by various pathogens remain a critical challenge for rainbow trout aquaculture, as they significantly affect fish health and production efficiency (Duman et al., 2025; Vervelacis et al., 2026). Breeding for pathogen resistance traits is particularly challenging given the high diversity of pathogens and the complexity of the fish immune system, which involves innate and adaptive responses (Makesh and Rajendran, 2022). Selective breeding leverages polygenic adaptation across mechanisms, unlike potentially limiting marker-assisted selection for single pathways. Consequently, identifying favourable QTLs or genes associated with resistance traits is essential for developing effective breeding programs to mitigate the risk and burden of infectious diseases (Robinson et al., 2023). Hence, several studies on rainbow trout have primarily focused on the identification of QTLs for disease resistance, identified through genetic linkage maps associated with traits such as time to death, or the binary survival/mortality (Liu et al., 2015; Palti et al., 2015; Vallejo et al., 2014). However, variability between studies and environmental conditions hinders the identification of robust QTL and their precise localization (Fraslin et al., 2020). MQTL analysis integrates data from multiple populations to identify robust genomic loci involved in a trait of interest, improving resolution and reliability (Elaswad and Dunham, 2018; Yáñez et al., 2023). To date, no comprehensive analysis combining QTLs for pathogen resistance is available in rainbow trout. Our study integrates 145 QTLs onto a new consensus genetic map resulting in 100 projected QTLs condensed into 26 MQTLs. The consensus map together with GWAS-validation enabled the precise localization of MQTLs associated with crucial pathogen resistance such as *F. psychrophilum* and IPNV, providing concise and accurate data to breeders and researchers for genomic selection strategies.

### 4.1 Genetic complexity and improvement precision

#### 4.1.1 MQTL analysis

Some studies have shown that resistance to different pathogens could be genetically correlated (Henryon et al., 2005; Evenhuis et al., 2015). Although an analysis with focus on a specific pathogen may reveal some of the associated loci, pooling correlated resistance traits measured within the same population can enhance the power to detect major QTL regions (Goffinet and Gerber, 2000). This meta-analysis approach operates under two different assumptions: 1) some of these traits share common underlying QTLs and 2) combining QTLs raises the likelihood of identifying genomic regions influencing multiple resistance traits simultaneously. The distribution of MQTLs across most chromosomes, each involving multiple loci with moderate effects, suggests to a wide extent a polygenic nature of pathogen resistance. This pattern is consistent with genetic studies performed in other aquaculture species such as tilapia and Atlantic salmon, in which disease resistance is rarely determined by a single major locus but rather arises from the complex interplay of numerous genes of minor effect (Houston et al., 2020). The widespread chromosomal distribution and moderate average PVE of MQTLs emphasize the importance of genetic improvement approaches that capture their polygenic architecture effectively to avoid information loss and overestimation of individual QTL effects (Nguyen, 2024; Tsai et al., 2015). The second assumption was supported by a significant reduction in CI following MQTL analysis. Across the genome, CI lengths of projected QTLs were reduced by 50.7% on average, corresponding to a 2.03-fold decrease compared with the original QTLs, which is in line with observations in MQTLs studies in plants (Sharma et al., 2024). The magnitude of the observed reduction, although significant, is more conservative than those reported in agriculture crops, which range from 5 to 6-fold reductions (Khojasteh et al., 2024; Kumar et al., 2023; Sandhu et al., 2021). This difference is likely due to the relatively early stage of genomics in rainbow trout compared to model plant species, where decades of intensive mapping have generated hundreds of QTLs (Rani et al., 2023).

All MQTLs were associated with resistance to *F. psychrophilum* since there were more studies on bacterial cold-water disease, highlighting its economical and epidemiological importance in aquaculture (Mathiessen et al., 2023; Wiegertjes and Elks, 2022). In general, MQTLs on chromosomes 3, 8 and 12 exhibited the highest number of MQTLs, potentially representing genomic hotspots for pathogen resistance in rainbow trout. This pattern aligns with the complex nature of immune traits observed in aquaculture species and emphasizes the importance of integrating data across multiple independent studies to refine key genomic regions (Houston et al., 2020; Yáñez et al., 2023).

The large number of initial QTLs in each MQTL and the significant reduction of CI, along with multiple QTLs from different studies co-locating at the same position, make that MQTL as one of the most notable regions associated with the favourable trait (Yang *et al*. 2021a). MQTL 3.1 represents the aforementioned pattern as stands out as the most promising region as it is associated with resistance to both *F. psychrophilum* and IPNV, with the largest CI reduction and highest number of projected QTLs from independent studies. This convergence across datasets underscores its relevance for pathogen-resistance genetics in rainbow trout.

#### 4.1.2 GWAS studies

GWAS is a compelling approach for exploring quantitative traits with higher resolution (Yáñez et al., 2023), which was used in rainbow trout to identify numerous genetic markers associated with pathogen-resistance (Calboli et al., 2022; D’Ambrosio et al., 2025; Palti et al., 2024; Silva et al., 2019). We combined available GWAS studies for disease resistance in rainbow trout showing the genome-wide distribution of significant markers for several pathogens. In line with the MQTL analysis, this map highlights the complex and polygenic nature of disease resistance traits, where multiple loci jointly influence resistance / susceptibility to diverse pathogens.

To enhance the precision of the study, significant genomic regions were defined by employing rigorous criteria requiring validation from multiple GWAS sources and considered effect sizes and consistency, which identified several regions associated with disease resistance on chromosomes 3 and 21. Chromosome 3 contains two loci linked to resistance against IHNV - *F. psychrophilum* and *F. columnare* - *F. psychrophilum*, respectively, while chromosome 21 harbours two regions related to resistance against *V. anguillarum* - *F. columnare* and IHNV - IPNV. These sub-megabase regions likely have pleiotropic effects and contain genes involved in immune defence. For instance, on chromosome 3, region flanking the SNP with the lowest p-value for the QTL associated with resistance to *F. columnare* (and *F. psychrophilum*) harbours two genes linked to proinflammatory cytokine responses (*tgf-beta2* and *il-1r1*) (Fraslin et al., 2022). On chromosome 21, the top SNP associated with resistance to IHNV-IPNV was located in a region with the gene encoding *interleukin-8* (Rodríguez et al., 2019). The other region on chromosome 21, associated with resistance to *V. anguillarum* - *F. columnare*, co-localises with the protein phosphatase 1 regulatory subunit 12A and various segments encoding immunoglobulin light chain kappa (Karami et al., 2020). These represent promising targets for further functional investigations into their putative roles in immune function and disease resistance (Solovieff et al., 2013).

#### 4.1.3 GWAS-MQTL

GWAS data were used to validate our MQTLs. More than 60% of the MQTLs co-localized with independent GWAS studies, which indicated the coherence of the two approaches in pinpointing loci associated with disease resistance traits. The non-colocalized MQTLs might be the result of incomplete genetic diversity coverage, differences in genetic material, GWAS limitations to common variants, environmental influences or lack of physical position for certain traditional markers in the *O. mykiss* genome (Bilgrami et al., 2023; Sharma et al., 2024). Among the MQTLs validated through GWAS, those which fulfilled the criteria described in the material and methods (section 2.7) were designated as high-confidence MQTLs. Notably, four hcMQTLs (3.1, 6, 8.2 and 25) overlapped with multiple independent GWAS studies, confirming their biological relevance and the need for further investigation to elucidate their causal genetic architecture. The hcMQTL 3.1 was the most robust and reliable candidate, as it integrated the highest number of projected QTLs, having the narrowest CI and colocalization with multiple GWAS studies. This convergence suggests its potential for containing causal variants for pathogen resistance in rainbow trout, making it a high-confidence target for fine-mapping and functional validation (Benner et al., 2016; Giuffra et al., 2025). Although hcMQTL 6, hcMQTL 8.2 and hcMQTL 25 also showed high colocalization, respectively, their broader CI limits their applicability in breeding programs. While these regions should not be disregarded, their reduced genomic precision necessitates additional fine-mapping and resequencing efforts before implementation in breeding frameworks (Daware et al., 2017).

### 4.2 Candidate genes within the hcMQTLs and their association with pathogen resistance

Of the eight identified hcMQTLs, five harboured candidate genes. Mining for candidate genes within these hcMQTLs identified 11 unique genes associated with the immune response in rainbow trout, distributed across five chromosomes. Three immune-relevant genes were located on hcMQTL 3.1, the most noteworthy region for disease resistance: two sequences encoding immunoglobulin (Ig)-like domain-containing proteins and *map3k19. Map3k19*, a member of the MAP3K kinase family, plays a critical role in regulating type I interferon production and NF-κB signaling, contributing to antiviral immunity (Guan et al., 2023). Although there is no specific report on *map3k19* expression in rainbow trout, mammalian research showed its activation in the ERK and JNK pathways, acting as a novel cellular regulator (Hoang et al., 2020). In hcMQTL 6 both the receptor *mlcr1* and *irf8* gene found. *Mc1r* belongs to the GPCRs family and modulates tissue damage and repair by regulating melanogenesis and inflammation (Cal et al., 2017; Fierro-Castro et al., 2022). Meanwhile, the transcription factor *irf8* is essential for the development and maturation of myeloid lineage cells, thus playing a critical role in antigen processing and immune activation (Salem et al., 2020). *Irf8* upregulation has been demonstrated in response to viral and mitogenic stimulation in rainbow trout, and it is key to orchestrate efficient immune responses (Holland et al., 2010).

Candidate genes in hcMQTLs on chr. 8 were *cxlc8, ltb4r2b, sp1pr1*, *trim110* and *cactin*. Multiple *cxcl8* (interleukin 8) genes act as strong proinflammatory mediators, recruiting and activating neutrophils and other immune cells, and amplifying local inflammation (Hu et al., 2011; Bird & Tafalla, 2015; Hu et al., 2011). In rainbow trout, *cxcl8* orthologs are expressed broadly after stimulation with viral and bacterial mimics (Laing et al., 2002; Rebl et al., 2014), and are significantly induced in spleen during viral infections (Purcell et al., 2004; Tafalla et al., 2005) and in monocyte-like cells contributing to proinflammatory cascades (Montero et al., 2009). Additionally, two different receptors genes, *s1pr1* and *ltb4r2b,* were found on the same region. The first gene, *s1pr1*, also belongs to the GPCR family, meanwhile *ltb4r2b* is involved in tissue damage and repair by regulating macrophage migration (Ermis et al., 2024). Their co-localization in *F. psychrophilum*-associated MQTLs, a disease characterized by tissue necrosis, ulcers and erosion (Lee et al., 2023), highlights their potential role in controlling excessive inflammation and promoting tissue repair under pathological challenges. The *trim110* gene is an E3 ubiquitin ligase belonging to the TRIM family, supporting the selective polyubiquitination of key regulatory immune proteins (Yang et al., 2020). The function of *trim110* has not been studied in rainbow trout but an increase of *trim110* has been shown in carp exposed to Grass carp reovirus infection (Qin et al., 2021; Yang et al., 2020).

Finally, within the interval on chr. 12 two different genes, *prf1.3 and G-csf*, were found. Cytotoxic mediators like perforin (*prf1.3*) are involved in immune cell cytotoxicity and myeloid cell development, and they are synthesized by CD8+ T cells and natural killer cells (Thiery and Lieberman, 2014). In rainbow trout, differential expression of perforin genes in lymphoid organs highlighted their role in adaptive immunity (Athanasopoulou et al., 2009; Takizawa et al., 2011), although some studies suggested functional redundancy among perforin isoforms (García-Álvarez et al., 2024). The *G-csf* gene regulates myeloid progenitor proliferation, neutrophil generation and systemic immune response initiation (Panopoulos and Watowich, 2008). In *O. mykiss*, *csf3b2* is expressed in the spleen and it was upregulated following bacterial stimulation, indicating amplified inflammatory and lymphoid cells responses (Mengqun et al., 2025).

## 5. CONCLUSIONS

Infectious diseases affecting rainbow trout aquaculture remain as one of the major challenges the sustainability of the industry and a threat to global food security. Numerous studies have identified genomic regions associated with resistance to pathogens in this species, but heterogeneity in pathogen types, genetic markers, and trout lines has led to inconsistent findings. In this study, we constructed a consensus genetic map of rainbow trout, integrating diverse genetic markers, and validated the predicted MQTL regions using GWAS studies for pathogen resistance. Results from this work enhance our understanding of the genetic control of resistance to both bacterial and viral diseases in rainbow trout, offering new insights for breeding for disease resistance. The 53.29-55.88 Mb region on chromosome 3 emerged as an especially robust genomic region associated with disease resistance in both MQTL and GWAS studies. In addition, we identified putative candidate genes within hcMQTLs likely involved in immune function, making these regions suitable targets for selective breeding programs. This research provided a valuable resource for scientists aiming to develop new rainbow trout lines with enhanced resistance to bacterial and viral pathogens.

## Funding

This study is supported by postdoctoral programme of the Xunta de Galicia (Consellería de Cultura, Educación, Formación Profesional e Universidades) to R. Rodríguez-Vázquez (ED481B-2023-104).

## Credit Authorship contribution statement

R.R.V.: Conceptualization, methodology, software, formal analysis, investigation, data curation, visualization, funding acquisition, writing - original draft, writing - review & editing; M.K.G.: Conceptualization, methodology, investigation, writing - review & editing ; A.K: data curation, investigation, writing - review & editing; D.R.: methodology, investigation, supervision, resources, funding acquisition, project administration, writing - review & editing; K.B.: conceptualization, methodology, investigation, supervision, resources, project administration, writing - review & editing.

## Declaration of competing interest

The authors declare no conflict of interest

## Data availability

Data will be made available on request

## Notes

### Competing Interest Statement

The authors have declared no competing interest.

