## Supplementary material for "The genetic basis of resistance to pathogens in rainbow trout: a meta-QTL analysis": Fig. S1

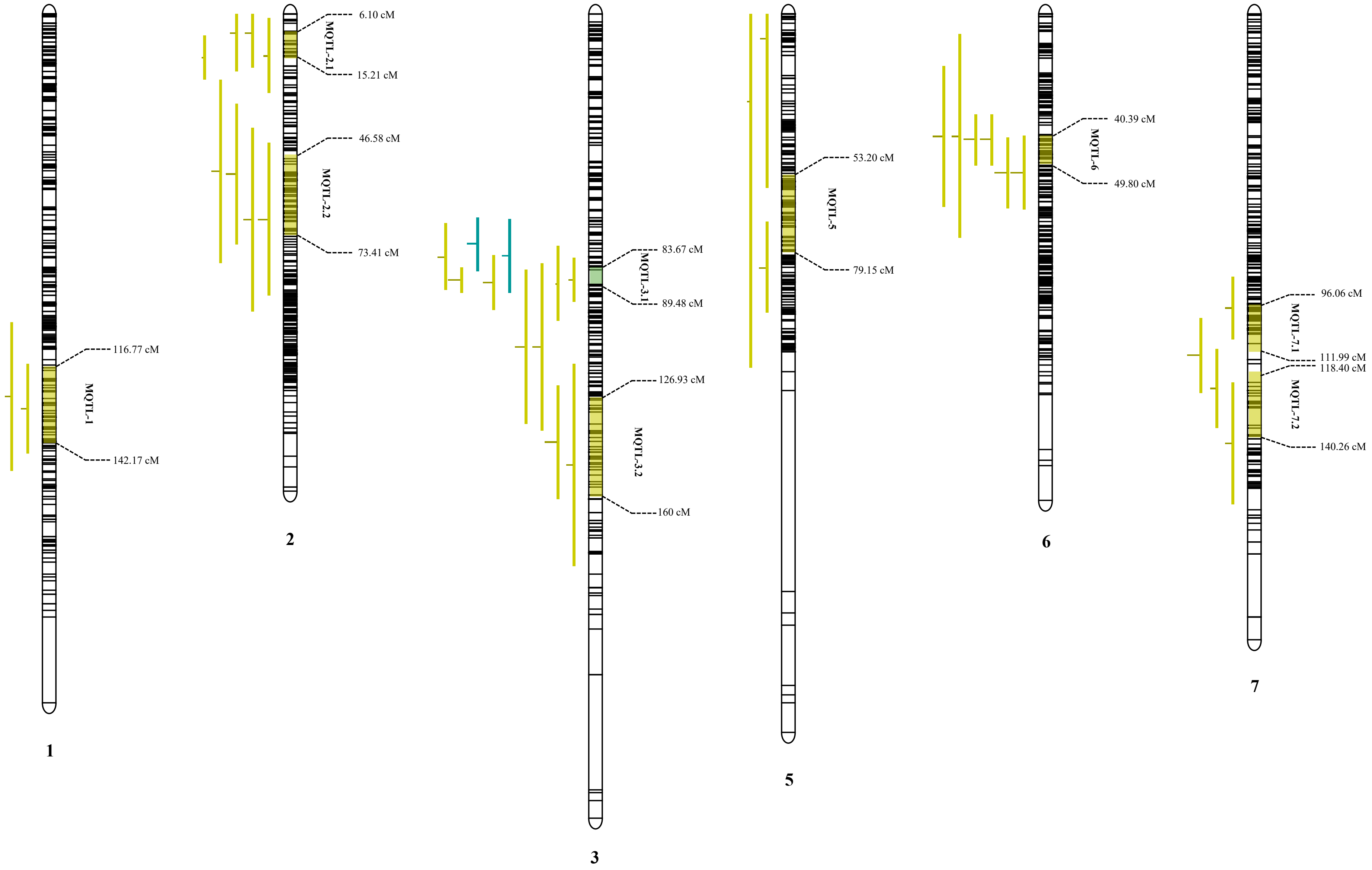

116.77 cM
MQTL-1
142.17 cM
1
6.10 cM
MQTL-2.1
15.21 cM
46.58 cM
MQTL-2.2
73.41 cM
2
83.67 cM
MQTL-3.1
89.48 cM
126.93 cM
MQTL-3.2
160 cM
3
53.20 cM
MQTL-5
79.15 cM
5
40.39 cM
MQTL-6
49.80 cM
6
96.06 cM
MQTL-7.1
111.99 cM
118.40 cM
MQTL-7.2
140.26 cM
7

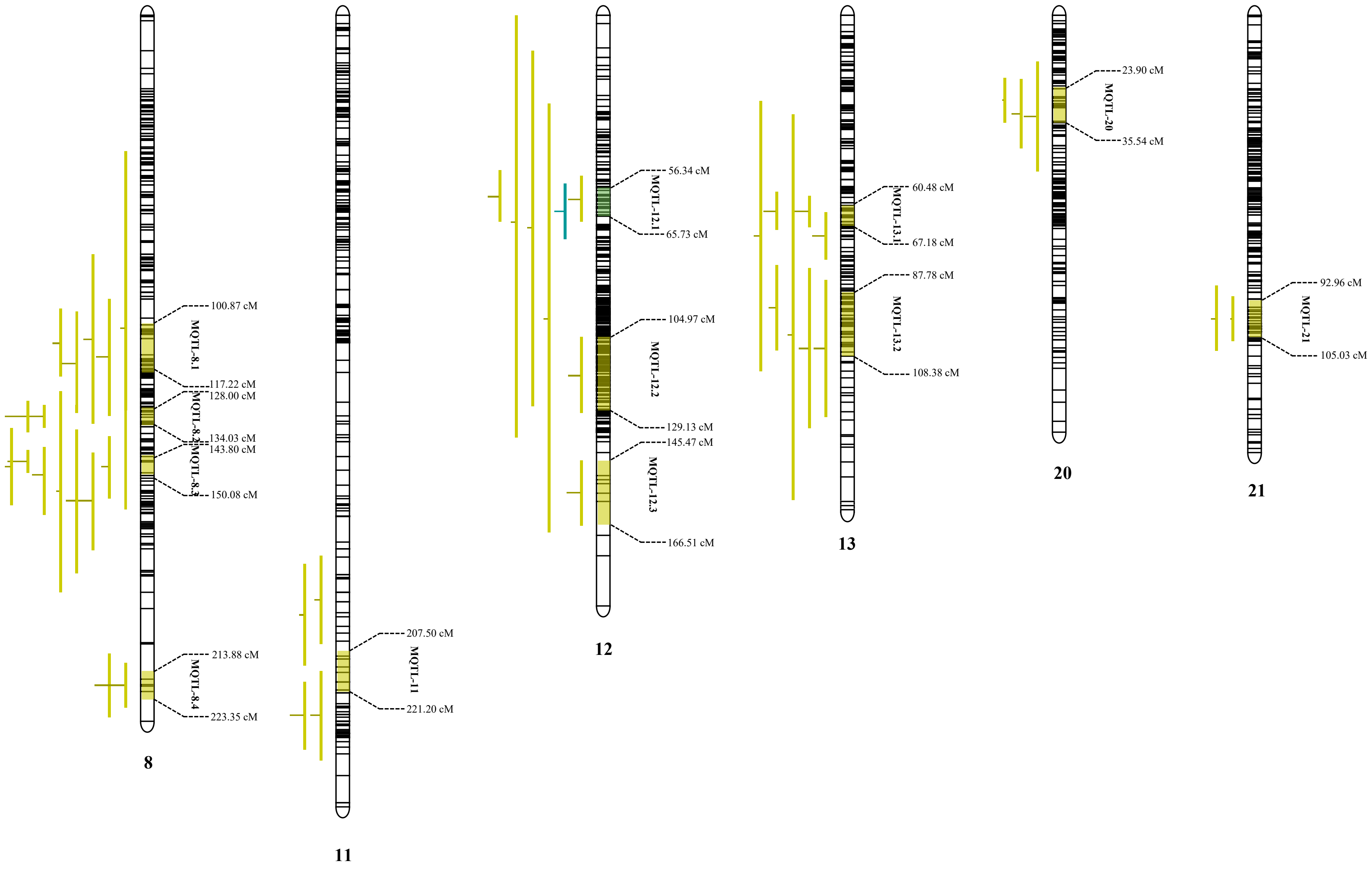

100.87 cM
MQTL-8.1
117.22 cM
128.00 cM
MQTL-8.2
134.03 cM
143.80 cM
MQTL-8.3
150.08 cM
213.88 cM
MQTL-8.4
223.35 cM
8
207.50 cM
MQTL-11
221.20 cM
11
56.34 cM
MQTL-12.1
65.73 cM
104.97 cM
MQTL-12.2
129.13 cM
145.47 cM
MQTL-12.3
166.51 cM
12
60.48 cM
MQTL-13.1
67.18 cM
87.78 cM
MQTL-13.2
108.38 cM
13
23.90 cM
MQTL-20
35.54 cM
20
92.96 cM
MQTL-21
105.03 cM
21

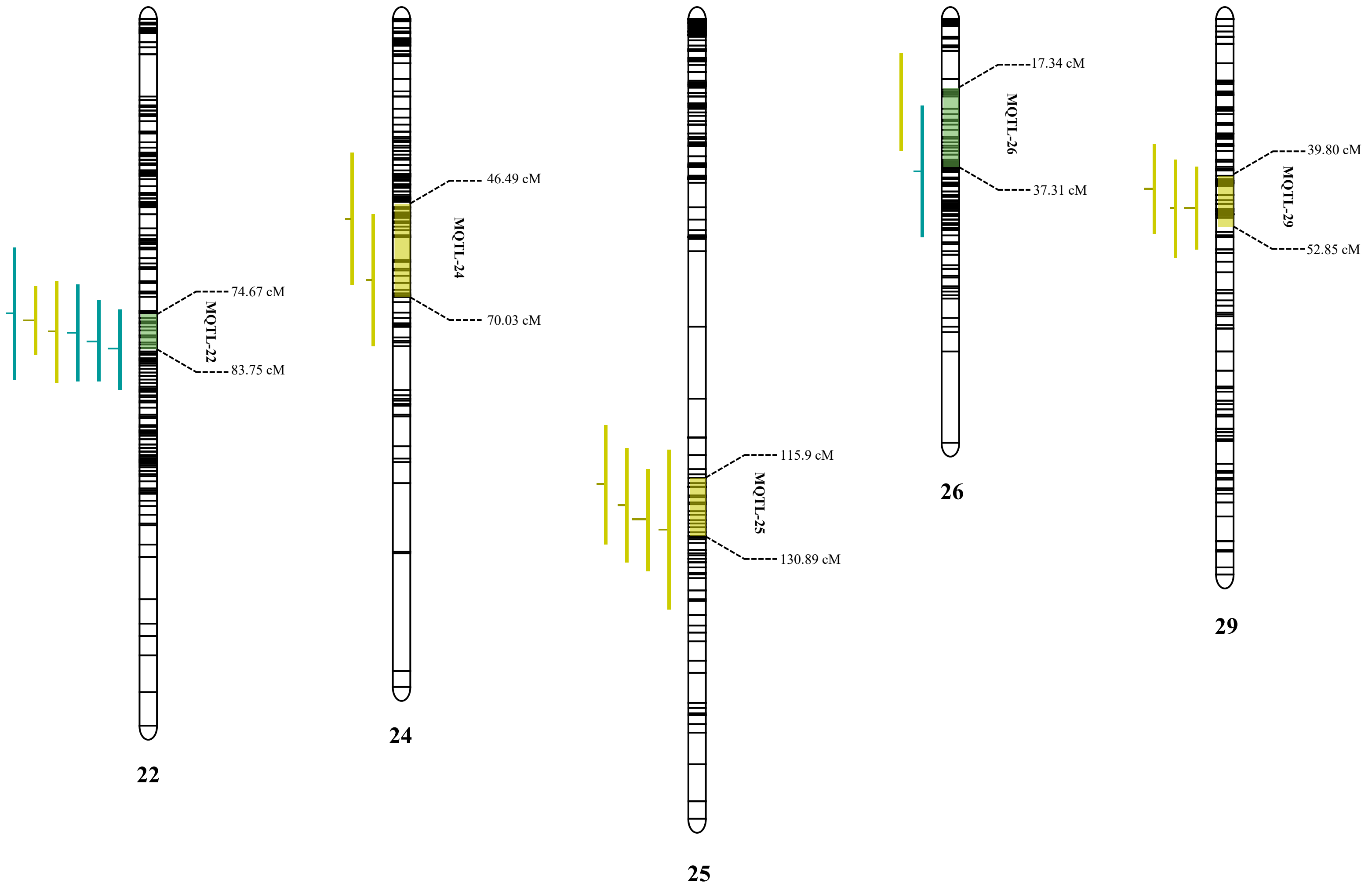


**Fig S1** . Schematic representation of the distribution of projected QTLs and MQTLs on rainbow trout chromosomes, associated with pathogen resistance traits. Bars on the left side of each chromosome indicate QTLs associated with *F. psychrophilum* resistance traits (green bars) and IPNV resistance traits (blue bars). Black bars within the chromosomes represent marker density. The chromosome map shows the MQTLs identified in this study and their co-location: light green indicates MQTLs formed by only one type of pathogen resistance QTL, while dark green indicates MQTLs formed by two different types of pathogen resistance QTLs.
